# Mutation Reporter: Protein-Level Identification of Single and Compound Mutations in NGS Data

**DOI:** 10.1101/2024.05.30.595736

**Authors:** Mikaela Teodoro, Rosana V. das Chagas, José A. Yunes, Natacha A. Migita, João Meidanis

## Abstract

**Summary:** Next-generation sequencing (NGS) has accelerated precision medicine by enabling simultaneous analysis of multiple genes and detection of low-frequency mutations. However, few open-source tools allow non-specialized users to transparently adjust quality parameters during mutation analysis. Mutation Reporter was developed to identify both single and compound amino acid alterations directly from raw fastq files of sequencing originated from RNA or exon sequences. The software provides full parameter control—including alignment e-value, minimum read length, minimum read depth, and minimum variant allele frequency (VAF).

**Availability and implementation:** Mutation Reporter is available to users through a free GNU software license and can be accessed on GitHub (https://github.com/meidanis-lab/mutation-reporter) and as a Code Ocean capsule (https://codeocean.com/capsule/0121109/tree).

## 1 Introduction

Next-Generation Sequencing (NGS) has transformed genomic research and precision medicine by enabling comprehensive analysis of multiple genes simultaneously. This high-throughput capability allows detection of mutations that may be implicated in treatment resistance, relapse, or poor prognosis in cancer and other genetic diseases [1, 2, 3]. The ability to identify such alterations underscores the clinical significance of NGS in therapeutic decision-making and disease monitoring.

However, the interpretation of NGS data remains a bottleneck for many research and diagnostic laboratories. Most available software packages for mutation analysis are designed for users with advanced bioinformatics expertise, requiring command-line operations, complex parameter adjustments, and familiarity with sequence alignment and variant-calling pipelines [4, 5]. As a result, laboratories without dedicated computational support often rely on commercial or semi-automated tools that lack transparency, flexibility, or detailed parameter control, which can compromise reproducibility and diagnostic accuracy.

An additional challenge arises when determining whether two or more mutations occur within the same DNA molecule — a configuration known as a compound mutation. In this context, compound mutations represent in cis events, in which double or multiple mutations occur on the same allele or sequencing fragment [6, 7]. This contrasts with polyclonal mutations, where different mutations originate from independent subclones or distinct alleles (in trans) within a cell population. Identifying such distinctions is clinically relevant, as compound mutations have been associated with poor therapeutic response and increased resistance to targeted therapies. In the case of BCR-ABL1, for example, differentiating compound from polyclonal mutations is essential because the presence of ≥ 2 mutations within the same molecule can confer markedly stronger resistance to tyrosine kinase inhibitors (TKIs), directly influencing therapeutic choice and clinical outcome [7, 8, 9]. Similarly, in EGFR, compound alterations have been associated with poor prognosis and reduced response to EGFR-targeted therapies [10, 11, 12].

Despite this clinical relevance, distinguishing compound from polyclonal events remains technically challenging. From an experimental perspective, many sequencing strategies do not generate fragments long enough to span two distant mutations, preventing direct assessment of whether they co-occur on the same molecule. Computationally, most existing analysis tools also lack mechanisms to verify co-occurrence across paired-end reads or to reconstruct haplotypes from short-read data. As a result, they typically treat variants independently, making it difficult to determine whether mutations belong to the same molecule or arise from separate subclonal populations

To address these limitations, we developed Mutation Reporter, an open-source tool that identifies amino-acid–altering individual and compound variants, including substitutions and in-frame insertions or deletions, directly from raw RNA or intron-free DNA FASTQ data. By performing protein-level alignment through BLASTX [13], the software enables intuitive detection of amino acid alterations while allowing users to adjust key analysis parameters such as alignment e-value, minimum alignment length, sequencing depth, and variant allele frequency (VAF) threshold. Mutation Reporter is openly available on GitHub and Code Ocean, ensuring transparency, accessibility, and reproducibility.

The main contributions of this work are summarized as follows:

- Development of an open-source and cross-platform software for mutation identification at the protein level.
- Introduction of a fast and parameterizable approach to detect both individual and compound mutations in paired-end reads.
- Demonstration of the software’s applicability in clinical sequencing data, highlighting its value for translational research.

The related work on mutation detection pipelines and comparative tools is reviewed in Section II. Section III describes the methodology and implementation details of *Mutation Reporter*. Section IV presents experimental setup, while Section V discusses implications and limitations. Section VI concludes the paper and outlines directions for future development.

## 2 Related Work

Numerous software tools have been developed for mutation detection in next-generation sequencing (NGS) data. Among these, VarScan [4] and GATK [5] are widely used in genomic studies, operating at the nucleotide level to identify single-nucleotide variants (SNVs) and small insertions or deletions. While these tools are robust and flexible, they require extensive preprocessing steps such as sequence alignment, variant filtering, and parameter tuning, which can be challenging for non-specialized users. Furthermore, their analysis is restricted to the nucleotide level, providing no direct information about the corresponding amino acid changes or potential compound events.

In the context of transcriptomic data, RNAMut [3] was introduced as a pipeline capable of identifying somatic mutations directly from RNA-sequencing reads. Unlike conventional variant callers, RNAMut performs comparisons at the protein level, thereby allowing the user to visualize amino acid substitutions without requiring separate translation or annotation steps. However, the internal quality thresholds and filtering criteria used in RNAMut are not exposed to the user, which limits transparency and reproducibility. Additionally, the software does not provide functionality for detecting compound mutations, restricting the analysis to independent variant events.

AGEseq [14] was designed for analyzing mutations in amplicon sequencing experiments, offering read-level visualization and user-controlled filtering parameters. However, its analysis is performed at the nucleotide level, and amino acid changes must be manually inferred from the observed substitutions. Moreover, because AGEseq treats paired-end reads (R1 and R2) as independent sequences, it can only detect multiple mutations if they occur within the same read. This limitation prevents the identification of compound mutations that span across paired reads of the same sequencing fragment.

Despite these advances, a major gap persists in the availability of tools that combine ease of use, open access, explicit control of analysis parameters, and the ability to detect compound mutations across paired-end reads. Table 1 summarizes a comparison between existing approaches and the proposed *Mutation Reporter*.

**Table 1.**
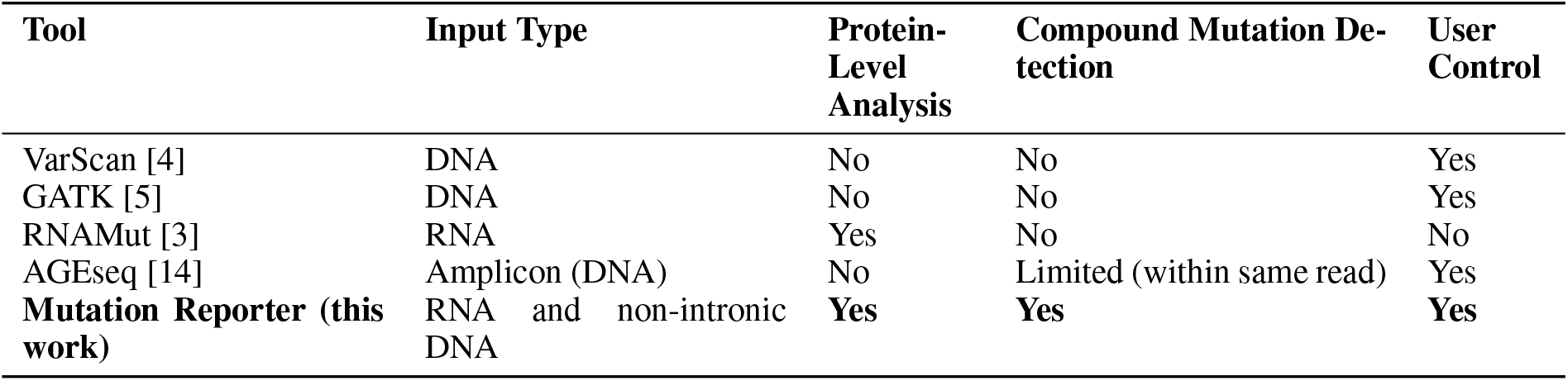
Comparison of mutation detection tools for NGS data.

## 3 Methodology and implementation

### 3.1 Overview

Mutation Reporter was designed as a modular, open-source software tool for the identification of amino-acid–altering variants, including both individual and compound mutations directly from raw RNA or intron-free DNA FASTQ data. The software combines usability, flexibility, and computational efficiency, enabling users without bioinformatics training to analyze sequencing results in a transparent and reproducible manner. Unlike conventional pipelines that rely on genome alignment and complex variant calling algorithms, Mutation Reporter performs a protein-level analysis using the BLASTX algorithm. This approach allows direct translation of DNA sequences into amino acid sequences, simplifying mutation identification and filtering based on user-defined quality criteria.

### 3.2 Software Architecture

Mutation Reporter follows a pipeline-based architecture organized into independent, sequential modules. Each module performs a well-defined task, receives input files from the previous step, and produces outputs for the next one. The pipeline is implemented using the *make* utility [15], which automatically manages dependencies and execution flow, improving reproducibility and maintainability.

The main components of the architecture are:

1. Preprocessing Module: Converts raw FASTQ reads (R1 and R2) into FASTA format suitable for BLAST analysis.
2. Merging Module: Combines paired-end reads into a single file, ensuring that reads from the same molecule share the same identifier.
3. Alignment Module: Executes BLASTX searches between query reads and the reference protein database provided by the user.
4. Mutation Extraction Module: Parses BLAST outputs, identifies mismatched amino acids, and records all detected mutations.
5. Reporting Module: Calculates variant allele frequencies (VAFs), determines single and compound mutations, and generates a summarized report.

This modular design facilitates parallel execution and debugging of the software. Our software was developed for Linux and it can also be executed seamlessly on macOS — which is Unix-based — and on Windows through the Windows Subsystem for Linux (WSL), a freely available Microsoft feature.

### 3.3 Mutation Detection Workflow

To prepare an analysis, users provide a FASTA file containing the reference peptide or protein sequences and a simple configuration file specifying analysis parameters such as maximum e-value, minimum alignment length, minimum identity, read depth, and minimum VAF threshold. Mutation Reporter then automatically builds a BLAST database from the input sequences using the *makeblastdb* utility, generating the necessary auxiliary files (.pdb, .phr, .pin, .pog, .pos, .pot, .psq, .ptf, and .pto) required for subsequent searches. This automation simplifies the setup process and eliminates the need for manual database preparation.

We selected BLASTX [13] for sequence alignment because it automatically translates nucleotide sequences in all six reading frames and aligns them against a protein reference. This feature enables direct identification of amino acid-level mutations without requiring additional translation or variant annotation steps, as would be necessary with other DNA-based tools such as BWA, Samtools, or VarScan.

BLAST can output results in multiple formats; we opted for XML because it provides detailed alignment statistics, sequence identity, and positional information. The XML output is parsed using the Bio.Blast.Record module from the Biopython package [16], allowing structured extraction of High-Scoring Segment Pairs (HSPs). Each HSP includes identity metrics, e-values, scores, and the start and end coordinates of aligned residues — data used to identify and quantify amino acid changes. All alignments are filtered according to user-defined parameters, including:

- **Maximum e-value:** threshold for acceptable alignment significance;
- **Minimum alignment length:** minimum number of amino acids required for a valid alignment;
- **Minimum percent identity:** minimum similarity between query and reference sequences;
- **Minimum read depth:** minimum number of reads supporting a mutation;
- **Minimum VAF:** minimum variant allele frequency to report.

The choice of minimum alignment length in Mutation Reporter is directly dependent on the sequencing design. Because the software operates on RNA-derived sequences or intron-free DNA amplicons, the parameter should be selected as a fraction of the expected translated fragment length. In practice, a minimum alignment length slightly below one-third of the read length is suitable for most amplicon-based applications. For example, for 150-bp reads, a threshold of approximately 49 amino acids ensures that only well-supported alignments are retained while filtering out reads with excessive mismatches. Similarly, the minimum read-depth parameter should be adjusted according to the depth achievable for the experimental design. Amplicon sequencing typically yields high coverage, allowing the use of stricter depth thresholds, whereas lower-depth experiments may require more permissive settings.

Filtered alignments are then analyzed to identify mismatched residues between query and reference sequences. Each mutation is annotated with its corresponding gene, amino acid position, and substitution type (e.g., p.R273H). The following subsections describe in detail the strategies used to quantify single mutations and to identify compound mutation events.

### 3.4 Individual Mutation Identification

The identification and quantification of individual amino acid–altering variants constitute the first analytical stage of Mutation Reporter. Through BLASTX alignment, each translated read is compared with the reference protein to detect amino acid–level mismatches, including substitutions as well as in-frame insertions or deletions. Within each valid HSP, Mutation Reporter scans the consensus alignment line represented by characters such as “+” or blank spaces (conservative or non conservative replacement). For each detected mutation, the program records the Illumina transcript ID, the reference and mutant amino acids, the affected gene, and the residue position. This structured annotation enables precise quantification of mutation frequency across all reads.

Detected variants are filtered using the quality parameters defined above, ensuring that only reliable calls are retained for downstream analysis.

To determine sequencing depth, defined as the total number of reads covering a specific position and required for VAF calculation, Mutation Reporter implements an indirect yet highly efficient scheme. Because sequencing depth varies along the peptide sequence, the most accurate way to determine it at a given position is to count all valid reads that include that position.

A naive approach would involve looping through every read to count how many alignments cover a given residue, which proved computationally expensive, requiring over one hour in preliminary tests. Instead, Mutation Reporter constructs a preprocessed data structure in *O*(*n*) time, where *n* is the number of valid hits, enabling subsequent depth queries in *O*(log *s*) time, where *s* represents the number of distinct start and end coordinates.

For a given position *x*, the depth is calculated as the difference between the number of reads that start before *x* and those that end before *x*. These values are obtained from cumulative count vectors that record the number of reads starting or ending at each coordinate. A binary search efficiently retrieves the cumulative values needed to compute depth at position *x* (Figure 1a).

**Figure 1.**
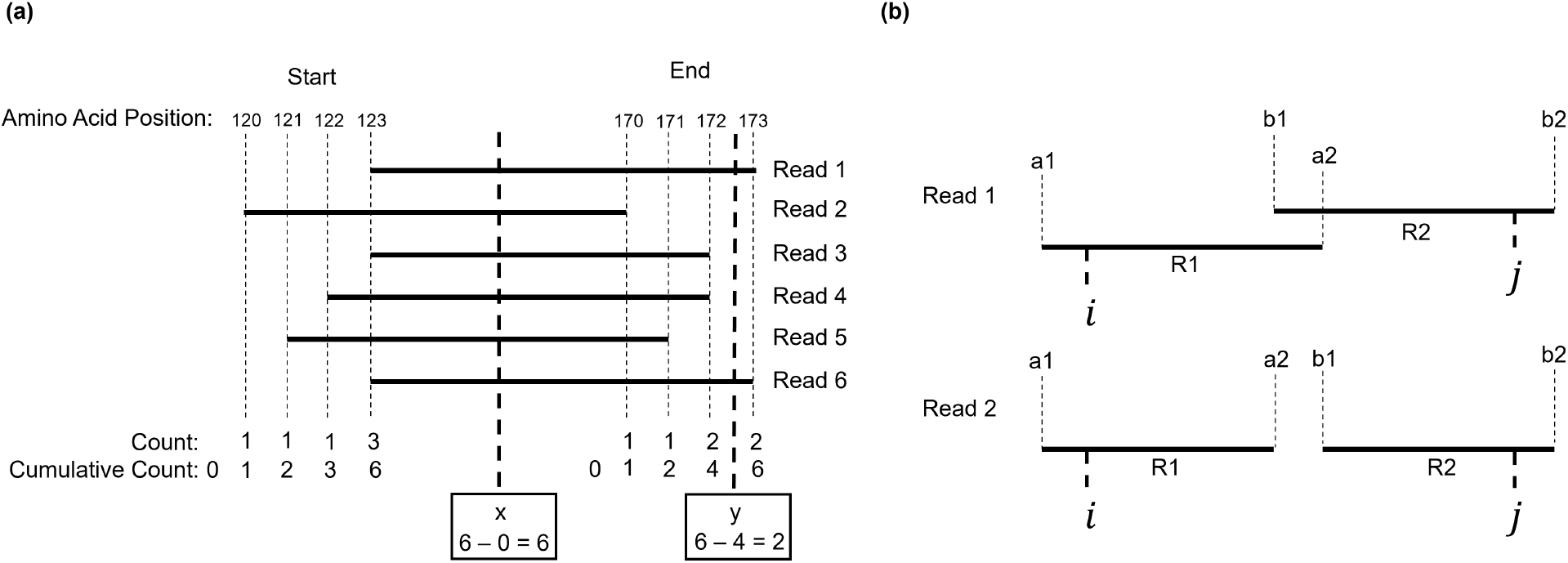
Preprocessed data structure for depth calculation. (a) Depth calculation for individual mutations, showing reads spanning amino acid positions, counts (number of reads starting at each position) and cumulative counts. Positions *x* and *y* indicate example depth calculations. (b) Two examples of transcript read intervals (R1, R2) and overlapping mutations (*i, j*), illustrating depth assignment.

Although more complex data structures such as interval trees [17] can also support interval-based queries, they are primarily designed for dynamic collections of intervals and would require significantly more implementation effort. Our simplified approach provides an effective balance between performance and simplicity.

This method is equally applicable to single-end and paired-end sequencing data. In single-end datasets, a compound mutation can be identified only when both mutations are covered within the same read. In paired-end data, co-occurrence is inferred also when both mutations are found within the same transcript ID, i.e., reads R1 and R2 belonging to the same DNA fragment. Once the sequencing depth at each position is determined, Mutation Reporter computes the VAF and reports only variants exceeding the user-defined frequency threshold.

The VAF for each mutation is calculated as:

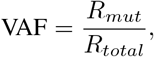

where *R*_*mut*_ is the number of reads containing the mutation and *R*_*total*_ is the total number of reads covering the corresponding position. Only variants exceeding the minimum VAF threshold are included in the final report.

This single-mutation framework forms the analytical foundation for the subsequent detection of compound events, described in the next subsection.

### 3.5 Compound Mutation Identification

Compound mutations are defined as the coexistence of two or more mutations within the same DNA molecule. Mutation Reporter identifies compound mutations by cross-referencing transcript IDs from paired-end FASTQ files. Reads sharing the same ID correspond to opposite ends of the same DNA fragment. For each pair of mutations *i* and *j*, Mutation Reporter calculates the compound VAF as the ratio between the number of transcripts carrying both mutations and the number of transcripts spanning the genomics positions of both. Formally:

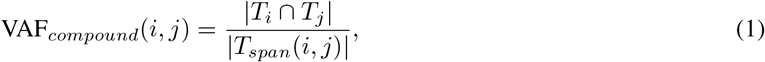

where *T*_*i*_ and *T*_*j*_ denote the sets of transcript IDs containing mutations *i* and *j*, respectively, and *T*_*span*_(*i, j*) represents the set of transcripts that span both positions. In paired-end sequencing, each read is represented by its genomic start and end coordinates: (*a*_1_, *a*_2_) for R1 and (*b*_1_, *b*_2_) for R2. These coordinates define the genomic interval covered by each read, and depending on the library preparation, the two reads may or may not overlap. A transcript is considered to span a given genomic position when that position falls within the interval covered by either R1 or R2 (Figure 1b).

To determine whether a transcript spans both mutations *i* and *j*, Mutation Reporter checks whether each mutation lies within either the R1 interval or the R2 interval. In other words, a transcript is counted as spanning the pair when position *i* is covered by R1 or R2, and position *j* is also covered by R1 or R2. This ensures that both genomic sites are physically present within the same DNA fragment represented by that transcript.

A compound mutation is detected when a transcript not only spans both positions but also contains both mutations simultaneously. This may occur when both variants are present within a single read (in single-end sequencing) or when one mutation lies in R1 and the other in R2 of the same paired-end fragment. Mutation Reporter calculates the compound VAF as the number of transcripts carrying both mutations divided by the number of transcripts that span both positions.

## 4 Experimental Setup

### 4.1 Datasets

To validate *Mutation Reporter*, we analyzed amplicon-based next-generation sequencing (NGS) data targeting the Ligand Binding Domain (LBD) of *RARA* and the B1/B2 boxes of *PML* within the *PML-RARA* fusion transcript. The dataset comprised 14 diagnosis samples and 34 samples collected at molecular relapse or in follow-up bone marrow aspirates with minimal residual disease (MRD) > 10^−4^. These samples were obtained from 15 pediatric patients diagnosed with Acute Promyelocytic Leukemia (APL) carrying the t(15;17) translocation and the *PML-RARA* fusion who showed a poor response to treatment according to the ICC APL Study One protocol. All of these patients were treated at the Boldrini Children’s Center (Campinas, Brazil). All procedures were approved by the institutional ethics committee under protocol CAAE 38962720.9.0000.5376.

Each sample consisted of paired-end FASTQ files (R1 and R2) generated using an Illumina MiSeq platform. The read length was 2 × 150 bp, with approximately 50,000–120,000 reads per file pair. Raw FASTQ files were preprocessed using Mutation Reporter’s internal conversion pipeline, and the analysis was carried out without prior alignment to a reference genome.

For the analysis of compound mutations (in cis), we used raw FASTQ files available in the European Nucleotide Archive (ENA). Viral RNA sequencing data generated on the Illumina MiSeq platform, with paired-end layout and amplicon-based library strategy, were analyzed. The samples correspond to SARS-CoV-2 viruses (Engl2 strain) cultured in vitro under selection with Remdesivir (RDV), from BioProject PRJNA692078 [18], which investigated the evolution of antiviral resistance. In total, we used 8 samples, resulting in 16 raw paired-end FASTQ files (R1 and R2) for the detection of compound mutations.

### 4.2 Validation Procedure

First, Mutation Reporter results of individual mutations were compared against those obtained using the *RNAMut* software. We selected RNAMut as the primary tool for comparison because it is the only publicly available software that performs mutation analysis at the protein level, directly reporting amino acid substitutions rather than nucleotide changes. Other widely used tools such as VarScan, GATK, and AGEseq operate exclusively at the nucleotide level, reporting variants in terms of base substitutions, which prevents direct comparison with Mutation Reporter’s amino acid-based output.

Variants identified exclusively by Mutation Reporter or RNAMut were visually inspected using the *Integrative Genomics Viewer (IGV)* [19] to confirm the presence of true mutations and to assess potential differences in VAF estimation. Discrepancies between the two methods were analyzed to identify sources of divergence, such as read mapping strategy, quality filtering, or minimum VAF thresholds.

The validation also assessed Mutation Reporter’s ability to detect compound mutations across paired-end reads, a feature not implemented in RNAMut. In this case, we assessed the presence of compound mutations by visual inspection using IGV.

The following parameters were applied in all Mutation Reporter executions:

- maximum e-value = 10^*−*5^;
- minimum alignment length = 49 amino acids;
- minimum percent identity = 90;
- minimum read depth = 500;
- minimum VAF = 2.

### 4.3 Performance Evaluation

To evaluate computational performance, a subset of datasets was selected to test the relationship between FASTQ file size and execution time. Specifically, 15 ABL1 samples (single-gene analysis) and 11 PML-RARA samples were processed on Machine 1, while an additional 21 PML-RARA fusion datasets—comprising two genes to be analyzed—were executed on Machine 2:

- **Machine 1:** Desktop computer equipped with an Intel^®^ Core™ i7-2600 CPU (8 cores), 8 GB RAM, running Ubuntu 22.04 LTS.
- **Machine 2:** Workstation equipped with an Intel^®^ Xeon™ E-2224G CPU (4 cores), 16 GB RAM, running Ubuntu 22.04 LTS.

These datasets covered a broad range of input sizes, enabling assessment of runtime scalability across different experimental configurations.

## 5 Results and Discussion

### 5.1 Validation of Mutation Reporter Outputs

For the comparative analyses, only mutation types detectable by both Mutation Reporter and RNAMut were considered. Single-nucleotide insertions or deletions, dinucleotide substitutions, and other unsupported variants were therefore excluded. A comprehensive list of all mutations identified by each tool is available in the “PML–RARA” section of Supplementary Table 1. Across 48 analyzed PML-RARA samples from 15 patients, four showed no detectable variants (VAF *≥* 2%) in either software. Among the remaining 44 samples, 160 mutations were identified by Mutation Reporter and 102 by RNAMut, with 66 shared between both tools (Supplementary 1). Of the 94 unique to Mutation Reporter, 64 presented VAF below 3%, and only one (p.S287L, RARA LBD) exceeded 8%, with a VAF of 99.45% — consistent with 99.62% in IGV. The absence of this mutation in RNAMut remains unexplained. Conversely, among the 36 mutations uniquely reported by RNAMut, 21 had VAF below 3%, and only one (Q245R) exceeded 8%; however, this variant was not supported by IGV inspection, which showed only 29 reads covering the region, compared with 10 in RNAMut. Since this value does not reach the 500-read threshold defined in Mutation Reporter, the tool appropriately excludes the mutation.

Figure 2 shows the comparison of VAFs for shared mutations between both tools (≥ 2%). Mutations detected exclusively by one of the softwares are not shown in the figure. Except for four evident outliers, points closely followed the identity line (x = y), indicating strong concordance between the two tools.

**Figure 2.**
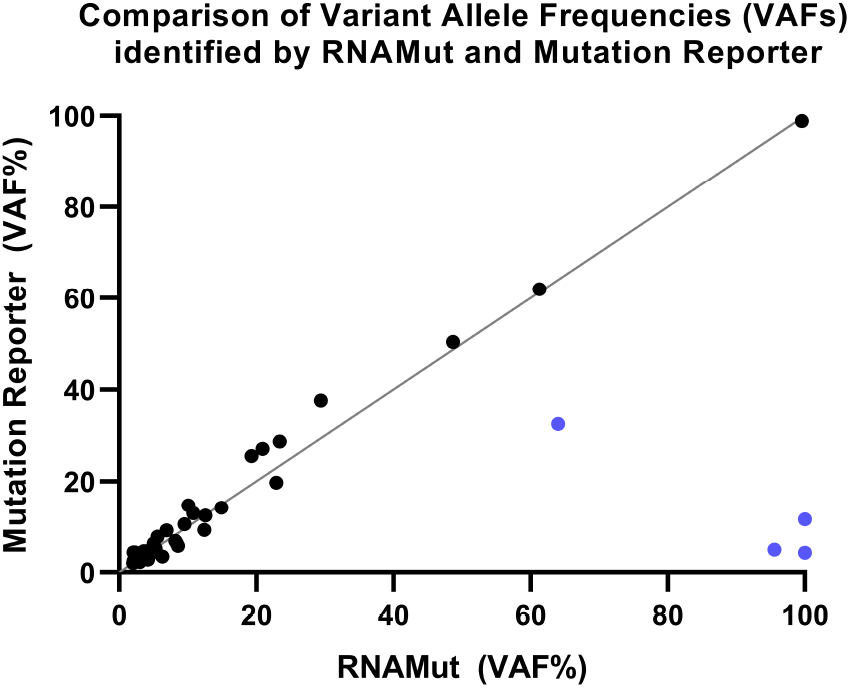
Comparison of variant allele frequencies (VAFs) identified by RNAMut and Mutation Reporter across 44 APL samples. A gray dashed line indicates the identity line (*y* = *x*), representing perfect agreement between both methods. Blue points highlight outlier mutations, which are discussed in Section V. Parameters: *e*-value = 10^*−*5^, minimum alignment length = 49, identity = 90%, depth = 500, and VAF *≥* 2%.

The four outliers corresponded to L218H (PML), I332T (RARA), M243T (PML), and T244A (PML), the last three in the same sample (S41, Supplementary 1). RNAMut reported markedly higher VAFs (64%, 95.6%, 100%, 100%) compared to Mutation Reporter (32.6%, 4.98%, 11.6%, 4.3%). In IGV, the actual VAFs were closer to Mutation Reporter’s (12% and 5% for M243T and T244A), with 717 reads covering the positions, while RNAMut showed much fewer total reads (71 and 21) and all carrying the variant, inflating the VAF. Similarly, I332T in RARA appeared with 5% in RNAMut and 5% in IGV, but Mutation Reporter had 1165 reads versus 45 in RNAMut. The L218H variant showed 61% in IGV, aligning more closely with RNAMut (61%) than with Mutation Reporter (33%). Overall, RNAMut appears to underestimate wild-type reads in several loci, leading to overestimated VAFs for those outliers.

### 5.2 Compound Mutation Analysis

For the PML-RARA fusion transcript, all compound mutations identified showed VAFs below 7%. As each event represents the proportion of transcripts carrying both substitutions relative to those covering their positions, mutations located in regions not co-covered by the same transcript were not identified as compound, reflecting the tool’s correct handling of read pairing.

To further validate this functionality, we analyzed RNA-seq data from severe acute respiratory syndrome coronavirus 2 (SARS-CoV-2), focusing on the S gene, which encodes the Spike glycoprotein. Eight samples were analyzed, revealing several co-occurring Spike mutations, with compound VAFs ranging from 0.07% to 99.71% (Supplementary 1). Figure 3 displays both the individual mutation VAFs (blue intensity) and their compound relationships (red lines), illustrating clonal complexity across samples.

**Figure 3.**
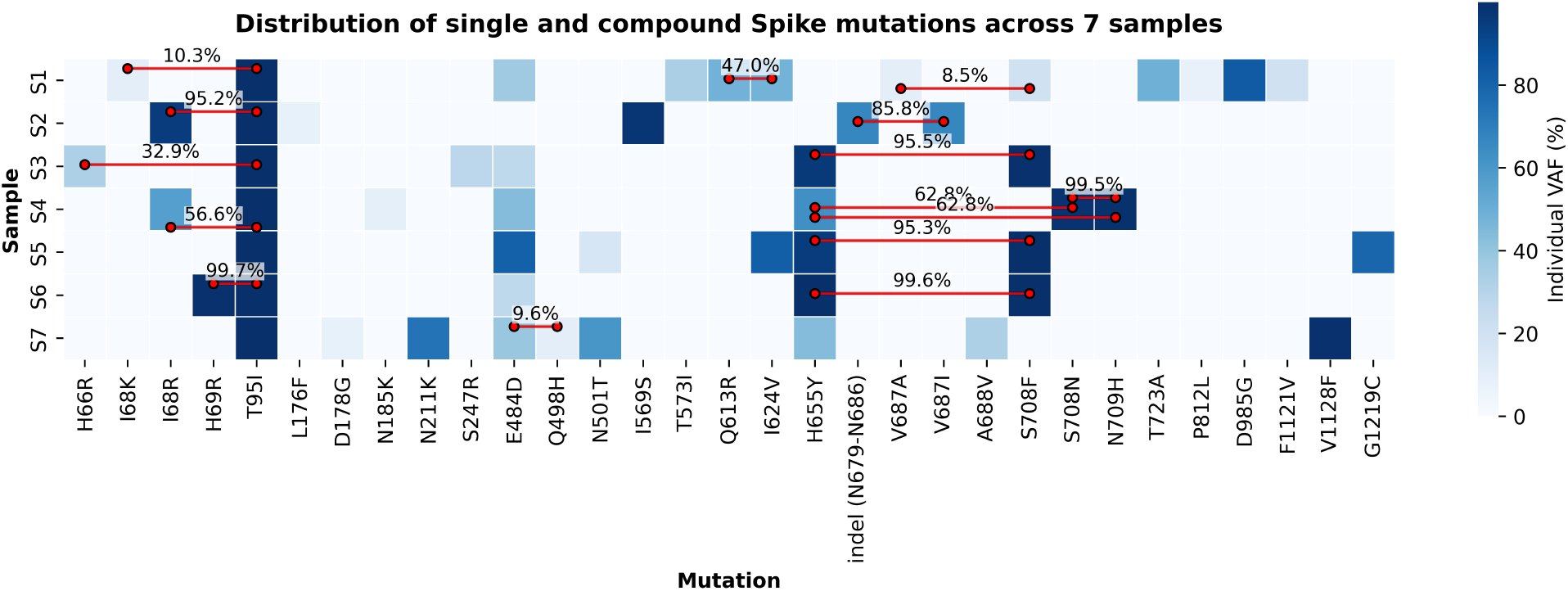
Distribution of single and compound Spike mutations across 7 DNA samples. The heatmap shows the variant allele frequencies (VAFs) of individual mutations (blue scale). Red lines connect mutation pairs detected as compound variants within the same sequencing reads, with labels indicating the compound VAF (%). Only single and compound events with VAF greater than 7% are displayed.

The individual mutations detected by Mutation Reporter corresponded precisely to those reported in the referenced study [20]. Notably, for mutation pairs in which one variant exhibits a VAF near 100%, that is, it is present in almost all reads, the compound VAF closely matches the individual VAF of the second mutation. This occurs because every read containing the lower-frequency mutation must necessarily also contain the nearly fixed mutation.

It is worth noting that there may be more compound mutations that were not reported due to the amplicon design. That is, only mutations present in the same transcript, according to the amplicon design and the primers used to amplify it, were reported. This allows for the planning of strategies to verify whether two mutations are compound or not. This analysis highlights Mutation Reporter’s potential for identifying variants occurring in combination. Such information may help reclassify currently uncharacterized variants as potentially pathogenic when found co-occurring with known driver mutations, an important step toward understanding mechanisms of tumor evolution and therapy resistance.

### 5.3 Performance Discussion

For datasets up to 50 MB (combined R1+R2), the full analysis was completed within 20 minutes on Machine 1. Larger datasets (65–135 MB) required approximately 60–90 minutes on Machine 2 (Figure 4).

**Figure 4.**
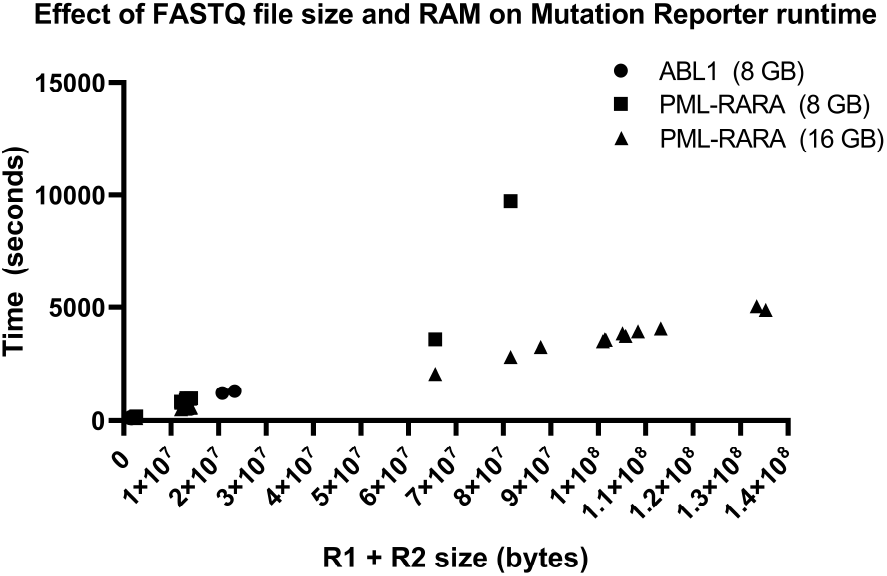
Runtime performance of Mutation Reporter for ABL1 and PML-RARA datasets under different memory conditions. Dots represent ABL1 samples (< 25 MB, ≤ 25 min), squares correspond to PML-RARA datasets (< 80 MB, ≤ 1 h on 8 GB RAM), and triangles denote larger datasets (< 135 MB, ≤ 1.5 h on 16 GB RAM).

In all cases, memory usage remained below 4 GB, confirming that the tool can be efficiently executed on standard laboratory hardware without specialized computational infrastructure. These results demonstrate the method’s scalability and suitability for integration into translational and diagnostic workflows.

Mutation Reporter exhibited near-linear scalability with respect to input size, confirming the efficiency of its modular design. Even when processing large paired-end datasets (¿100 MB combined), total runtime remained below 90 minutes on standard laboratory hardware. This level of performance ensures that the tool can be incorporated into real-world clinical and research workflows without requiring high-performance computing infrastructure.

From a usability perspective, Mutation Reporter’s open-source implementation and parameter transparency make it adaptable to a wide range of experimental contexts. The command-line interface allows reproducible execution via configuration files, and its modular Makefile architecture supports parallelization and pipeline integration with existing bioinformatics environments.

### 5.4 Current Limitations

The current version of Mutation Reporter is optimized for identifying point mutations and in-frame indels that alter amino acid sequences. Frameshift variants, large insertions or deletions, and structural rearrangements (e.g., gene fusions or tandem duplications) are not yet supported. Additionally, while the tool operates efficiently on standard laboratory hardware, very large datasets (¿150 MB combined FASTQ) may require extended runtime or higher memory resources. Additionally, although the BLASTX-based approach simplifies alignment, it may be less efficient for whole-transcriptome datasets due to the increased search space. Integration with variant databases in future versions would facilitate interpretation of mutation patterns in larger clinical cohorts.

## 6 Conclusion

This work presented *Mutation Reporter*, an open-source and platform-independent software designed for the identification of both single and compound mutations from next-generation sequencing (NGS) data. The proposed approach employs protein-level analysis through *BLASTX* alignment, allowing direct detection of amino acid substitutions and enabling transparent control over analytical parameters such as e-value, alignment length, sequencing depth, and variant allele frequency (VAF).

Validation using clinical datasets from pediatric acute promyelocytic leukemia (APL) samples demonstrated that Mutation Reporter accurately identifies known mutations and detects additional low-frequency variants overlooked by existing tools such as RNAMut. Furthermore, its ability to determine compound mutations across paired-end reads represents an important advancement for investigating co-occurring variants within the same molecule—an aspect often linked to clonal selection and therapeutic resistance.

Mutation Reporter operates efficiently on standard laboratory hardware, with near-linear time scalability and minimal memory usage. Its modular design and open-source distribution make it adaptable for diverse research and diagnostic environments, providing reproducibility, parameter transparency, and accessibility for users with varying levels of computational expertise.

Future work will focus on expanding variant detection capabilities to include frameshift mutations and structural rearrangements, as well as integrating a graphical user interface (GUI) for simplified use in translational and clinical contexts. By combining flexibility, accuracy, and ease of use, Mutation Reporter contributes to the growing need for transparent and reproducible bioinformatics tools in precision medicine.

## 8 Author contributions statement

MT implemented the final version of the software, wrote the manual, and conducted the experiments, RVC. implemented an initial version of the software, JAY. supervised the NGS of PML-RARA and reviewed the manuscript, NAM. conceived the idea for the software and reviewed the manuscript, JM supervised the implementations, helped with the writing of both the manual and the article, and organized the code repository.

## 9 Acknowledgments

The authors would like to thank the Centro Infantil Boldrini for providing access to sequencing data and the computational facilities used in this study.

